# Fingolimod acutely facilitates the activation of TRKB

**DOI:** 10.64898/2026.02.27.707919

**Authors:** Cecilia Anna Brunello, José Pedro Araujo, Nina Seiffert, Katja Kaurinkoski, Plinio Casarotto, Caroline Biojone, Eero Castrén

## Abstract

Fingolimod (FNG) is a sphingosine-1-phosphate receptor agonist currently prescribed for the treatment of remitting-relapsing multiple sclerosis. However, an increasing body of evidence indicates that FNG has a variety of other effects on the central nervous system, making it a good candidate to target other brain disorders that display loss of neuronal cells and synaptic plasticity. FNG treatments induce production of brain-derived neurotrophic factor (BDNF), which promotes neuronal plasticity, arborization and survival via signaling through its cognate receptor TRKB. In this study we characterize the relationship between FNG and TRKB *in vitro* and *in vivo* following acute treatments. We found that FNG induces TRKB activation in primary neuronal cultures in a BDNF-dependent way, indicating a rapid effect of FNG. This effect is different from the one elicited by antidepressants and is likely mediated by modulation of plasma membrane properties, as the enhancement of fluoxetine binding and dimerization of the cholesterol-insensitive TRKB mutant Y433F mimic the effects of cholesterol. Moreover, acute FNG treatment normalizes the generalization of conditioned fear response seen in heterozygous BDNF null female mice without affecting the wild-type littermates. Taken together, our data indicate that FNG allosterically promotes TRKB signaling and thereby induces the increase in BDNF production, which mediates the therapeutic effects of the drug on neuronal plasticity.

## INTRODUCTION

Fingolimod (FNG, FTY720) is a sphingosine-1-phosphate receptors agonist with immunosuppression activity on lymphocytes (Mandala et al. 2002). The compound is currently prescribed to treat relapsing-remitting multiple sclerosis. However, accumulating evidence over the past decade aiming at further elucidating FNG mechanisms of action indicate that the drug has a much wider variety of effects on different cell types and in different organs. In particular, a solid effect of FNG treatment both *in vitro* and *in vivo* is the increased level of brain derived neurotrophic factor (BDNF) mRNA and expressed protein in neuronal cells, accompanied by increased spinogenesis and dendritic arborization (Deogracias et al. 2012; Di Pardo et al. 2014; Miguez et al. 2015; Patnaik et al. 2020). Moreover, in animal models of different diseases, FNG rescue motor and cognitive symptoms through increased neuronal plasticity and survival (Deogracias et al. 2012; Kartalou et al. 2020; Miguez et al. 2015; Naegelin et al. 2021; Vidal-Martinez et al. 2019).

BDNF is a neurotrophic factor that, through its cognate receptor neuronal receptor tyrosine kinase 2 (NTRK2, TRKB), modulates synaptic and neuronal plasticity in the brain in an activity-dependent way (Zagrebelsky and Korte 2014). Moreover, BDNF is implicated in the pathophysiology of several brain disorders (Dou et al. 2022). Reduced BDNF levels have been found in brains and serum of severely depressed patients (Dwivedi et al. 2003; Karege et al. 2005), as well as in patients affected by Alzheimer’s, Parkinson’s and Huntington’s disease (Buchman et al. 2016; Y. Huang, Huang, and Yun 2019; Zuccato et al. 2008). In addition, reduction in BDNF-TRKB signaling occurs during chronic stress response (Castrén and Monteggia 2021; Duman and Monteggia 2006; Smith et al. 1995). Taken together, this evidence suggests that imbalances in BDNF-TRKB signaling pathways affect brain functions at different levels and that maintenance or restoration of physiological synaptic plasticity might prevent or ameliorate pathological conditions that manifest with very different symptoms.

While the involvement of BDNF as a therapeutic target of FNG treatment is well established, the molecular mechanisms of BDNF upregulation and promotion of synaptic plasticity remain unclear. TRKB signaling modulates BDNF gene expression in a positive feedback manner (Tuvikene et al. 2016; Yasuda et al. 2007), possibly contributing to the increase of BDNF levels induced by chronic FNG treatment (Deogracias et al. 2012; Doi et al. 2013; Golan et al. 2019; Vidal-Martinez et al. 2019; Yu et al. 2023). Traditional and rapid acting antidepressants also upregulate BDNF production (B. Chen et al. 2001; Deyama and Duman 2020) and we recently demonstrated that they directly act on TRKB receptors as positive allosteric modulators of BDNF signaling (Casarotto et al. 2021). In particular, antidepressants interact with the cholesterol-sensing domain in the juxtamembrane portion of the transmembrane domain of TRKB (Cannarozzo et al. 2021; Casarotto et al. 2021; Fred et al. 2019), raising the possibility that FNG could also modulate the activity of TRKB in a similar fashion. In this study, we investigate the potential interaction between FNG and TRKB as an early event of FNG-induced plasticity.

## METHODS

### Animals

Female adult mice (C57BL/6J-000664, Jackson Labs, maintained in the Laboratory Animal Center of the University of Helsinki), 16-18 weeks old at the beginning of the experiment, carrying a deletion in one of the copies of the Bdnf gene (BDNF.het) or their wild-type littermates were used (Karpova et al. 2011; Laukkanen et al. 2021). The experiments were conducted according to international guidelines for animal experimentation and the County Administrative Board of Southern Finland (ESAVI/38503/2019, ESAVI/40845/2022).

### Drugs

Fingolimod hydrochloride (FNG, 2-Amino-2-[2-(4-octyl-phenyl)-ethyl]-propane-1,3-diol hydrochloride; FTY720, #SML0700, Sigma-Aldrich) was dissolved in DMSO for *in vitro* experiments and suspended in Tween20 2% in saline for *in vivo* administration (intraperitoneal, ip; 1 mg/kg volume). Pravastatin (Orion Pharma) was dissolved in water for oral administration. Fluoxetine (Bosche Scientific #H6995) was biotinylated with EZ Link NHS PEG kit (Thermo scientific). Biotinylated fluoxetine (bFLX), biotinylated BDNF (bBDNF; Alomone Labs, #B-250-B), BDNF (PeproTech, #450-02) and TRKB_FC chimera (R&D biosystems, 688-TK) were dissolved in PBS. PP1 (Sigma-Aldrich, #P0040) was dissolved in DMSO.

### Cell culture

E18 neuronal primary cortical cultures were prepared as previously described (Sahu et al. 2019). Neurons were maintained in neurobasal media (Invitrogen) supplemented with 2% v/v B27 (Invitrogen) and 1% v/v penicillin/streptomycin and L-glutamine (Lonza). Neurons (DIV 10) were pretreated with TRKB_FC (10 μM) or PBS vehicle for 15 min and further treated for 30 min with FNG (0.1, 1 or 10 μM in DMSO) or DMSO vehicle and then processed for ELISA.

Neuronal cell line N2A was used for PCA experiments, while HEK293T cell line was used for the binding assay. Both cell lines were maintained at +37 °C 5% CO_2_ in DMEM (Lonza) media supplemented with 10% v/v inactivated fetal bovine serum (Invitrogen), 1% v/v L-glutamine and penicillin/streptomycin (Lonza).

### ELISA

Detection of phosphorylated TRKB and interaction with PLCg1 was done with ELISA as previously described (Casarotto et al. 2021; Laukkanen et al. 2021). Briefly, cortical primary neurons were plated 250,000 cells/well on 24 well-plates and treated with FNG (10 μM, 30 min). Cells were washed once with PBS and lysed (20 mM Tris-HCl; 137 mM NaCl; 10% glycerol; 0.05 M NaF; 1% NP-40; 0.05 mM Na3VO4 with protease and phosphatase inhibitors (Sigma). Neuronal lysates were transferred on transparent 96 well-plates previously coated overnight at +4 °C with total TRKB antibody (R&D, #AF1494) 1:1000 in carbonate buffer and blocked for 2h with 2% BSA (Sigma) and PBS 0.1% Tween (PBS-T). The plate containing the lysates was transferred back to +4 °C overnight. Next day, the plate was washed with PBS-T and further incubated overnight at +4 °C with the biotinylated secondary antibody anti-phosphotyrosine (BioRad; #MCA2472B), anti-phospho Src family Tyr416 (Cell Signaling Technologies; #6943) or anti PLCg1 (Cell Signaling Technologies; #5690) 1:2000 in PBS-T. After a final washing step, the plate was incubated for 2h at room temperature with HRP-conjugated streptavidin (Thermo Scientific) 1:5000 in PBS-T. Chemiluminescence was measured with a Varioskan plate reader (Thermo Scientific) following the addition of ECL substrate (Thermo Scientific).

### Surface TRKB

Surface TRKB experiments were conducted as previously described (Casarotto et al. 2021; Fred et al. 2019). Briefly, primary cortical neurons were grown 60,000 cells/well in 96 well-plates and treated with FNG (0, 0.1, 1 and 10 μM, 30 min) before being washed with PBS and fixed with 4% paraformaldehyde (PFA) for 20 min at room temperature. PFA was removed and cells were washed three additional times with PBS, then incubated for 1h at room temperature in blocking buffer without detergents (5% nonfat dry milk, 5% Bovine Serum Albumin in PBS) and left overnight at +4 °C with the primary antibody anti total TRKB (R&D, #AF1494) in blocking buffer. The next day, the primary antibody was washed away, and cells were incubated with the IgG anti-goat HRP-conjugated antibody for 1h at room temperature. Finally, following 4x 10 min washes with PBS, ECL substrate was added and chemiluminescence was measured with a Varioskan plate reader (Thermo Scientific).

### Protein-fragment complementation assay (PCA)

PCA is based on a humanized *Gaussia princeps* luciferase system originally described in Prof. Michnick laboratory (Remy and Michnick 2006). Specifically, split and complementary portions of the luciferase were cloned on the C-terminus of the rat TRKB receptor in pcDNA3.1/zeo backbones. Interaction of the overexpressed proteins of interest, here two TRKB molecules, allows the reporter luciferase to refold in its original and functional conformation and to produce light as PCA signal, following the addition of its substrate native coelenterazine, in a proportional amount of interacting TRKB molecules at that given time.

PCA experiments were conducted as already described (Brunello, Yan, and Huttunen 2016; Merezhko et al. 2018). Briefly, 10.000 N2A cells/well were seeded in white-walled 96 well plates and transfected 24h after plating with the phGLuc-TRKB plasmids (110 ng of DNA per well) with Lipofectamine 2000 (Invitrogen) according to manufacturer’s instructions. Transfected cells were incubated for an additional 24h. FNG treatments were performed 30 min before measurements and culture media containing the treatments was changed to phenol red-free and serum-free DMEM (Invitrogen). PCA signal was measured as luminescence with a Varioskan plate reader (Thermo Scientific) immediately after the injection of the substrate for the *G. princeps* luciferase, native coelenterazine (25 μM, NanoLight Technology).

### Ligand binding assay

The cell-free binding assay was performed as previously described (Cannarozzo et al. 2021; Casarotto et al. 2021). Briefly, HEK293T cells transfected with TRKB.wt construct were lysed (see ELISA section for details) and transferred to transparent 96-wells plates (HB, Perking Elmer) previously coated overnight with TRKB antibody (1:1000 in carbonate buffer, +4°C) and blocked with 3% BSA in PBS for 2h at room temperature. The samples were incubated overnight on the shaker at +4°C. The following day, after 3x 10 min PBS washes, biotinylated BDNF (0, 2, 4, 6, 8, 10 ng/ml, Alomone Labs, #B-250-B) was added with or without FNG (0, 0.1, 1, 10 μM) for 1h at room temperature. For the competition binding assay, a fixed concentration of Fluoxetine (1 μM) was incubated with increasing concentration of FNG (0, 0.01, 0.1, 1, 10, 100 μM). After washing away the treatments, the samples were incubated with HRP-conjugated streptavidin (1:10000, Thermo Scientific) for 1h at room temperature. Luminescence was measured with a Varioskan plate reader (Thermo Scientific) after the addition of ECL substrate according to manufacturer’s instructions (Thermo Scientific).

### qPCR

N2a cells were tested for the endogenous expression of TRKB. As a control, cells were transfected with a TRKB.wt plasmid as described in the previous section. Primers for assessing levels of TRKB mRNA following FNG treatment were synthetized by Sigma-Aldrich. TRKB primers were: forward, GAAGGGAAGTCTGTGACCATTT-3′, reverse: 5′-GTGTGTGGCTTGTTTCATTCAT-3′. Beta actin was used as housekeeping gene: TGTCACCAACTGGGACGATA-3′, reverse: 5′-GGGGTGTTGAAGGTCTCAAA-3’. RNA isolation from lysates was performed with PureLink RNA Mini kit (Thermo Scientific), while reverse transcription of RNA was carried out with LunaScript RT SuperMix Kit (New England Biolabs), according to manufacturer’s instructions. SYBR Green Maxima kit (Thermo Scientific) and BioRad CFX Opus 96 PCR cycler were used to detect relative mRNA amounts. Analysis was performed using the 2^ΔΔCT^ method in Microsoft Excel.

### Contextual discrimination task

The paradigm used for contextual fear conditioning was adapted from a previous study (Laukkanen et al. 2021). We used 25 animals between BDNF.hets and their wild-type littermates, and 5 animals per group for the pravastatin cohort. Pravastatin (10mg/kg) was administered for 14 consecutive days in drinking water. The animals were anesthetized with isoflurane (2% for 4 min) and received either vehicle or FNG (1 mg/kg) by intraperitoneal injection twice during the experiment: 2h before the conditioning and 2h before the exposure to the unfamiliar context. The conditioning was performed in context A (transparent walls 23x23x35 cm with metal grid bottom) for a total of 10 minutes, including a short habituation, the foot-shock session (0.6 mA/2 seconds, at intervals of 30 seconds/1 minute) and the last 5 minutes of recording of the freezing behavior of the animals. After 24h, animals were placed in the unfamiliar context B (box of same dimensions of context A but with black walls and black sleek bottom, no foot shocks) for 5 minutes. After 24h, animals were returned to context A for 5 minutes, without receiving any shock. Animal freezing (measured in seconds) was recorded with the TSE software (Bad Homburg, Germany).

### Statistical Analysis

The data was analyzed in Prism Graphpad 8.1.1 by Student’s t-test (two tails), one-, two and three-way ANOVA or Kruskal-Wallis as non-parametric tests, followed by Šidák, Tukey and Dunn’s post hoc tests. Replica numbers as well as specific p-values are indicated in figure legends.

## RESULTS

### FNG promotes TRKB dimerization and signaling in vitro

While other studies have investigated the effects of FNG on BDNF and TrkB signaling after 24 h of administration (Deogracias et al, 2014; Patnaik et al, 2020), we have here focused on early effects of FNG on TrkB activity. We first assessed whether acute FNG treatments (0.1, 1, and 10 μM, 30 min) are able to induce TRKB homodimerization in N2A cells. These cells do not express endogenous TRKB (Supplementary Fig. 1A), however express small levels of BDNF (H. Chen, Lombès, and Le Menuet 2017; Haapasalo et al. 1999). FNG dose-dependently induced TRKB dimerization, with a significant effect at 10 μM (Fig. 1A). Similarly, when we tested by ELISA on cortical primary neurons whether TRKB was phosphorylated as a result of its activation, we found that FNG dose-dependently increases TRKB phosphorylation at the Tyr 816 site and this effect reached significance, more than threefold compared to the vehicle-treated control, at 10 μM (Fig. 1B). Consistently, the interaction between TRKB and its signaling partner PLCg1 was doubled at 10 μM FNG in cortical neurons (Fig. 1C), indicating that TRKB was activated, phosphorylated, and that PLCg1 signaling cascade was initiated. The effects on TRKB mimic the effects elicited by TRKB natural ligand, BDNF (Supplementary Fig. 1B, 1C. 1D, 1E).

**Figure 1.**
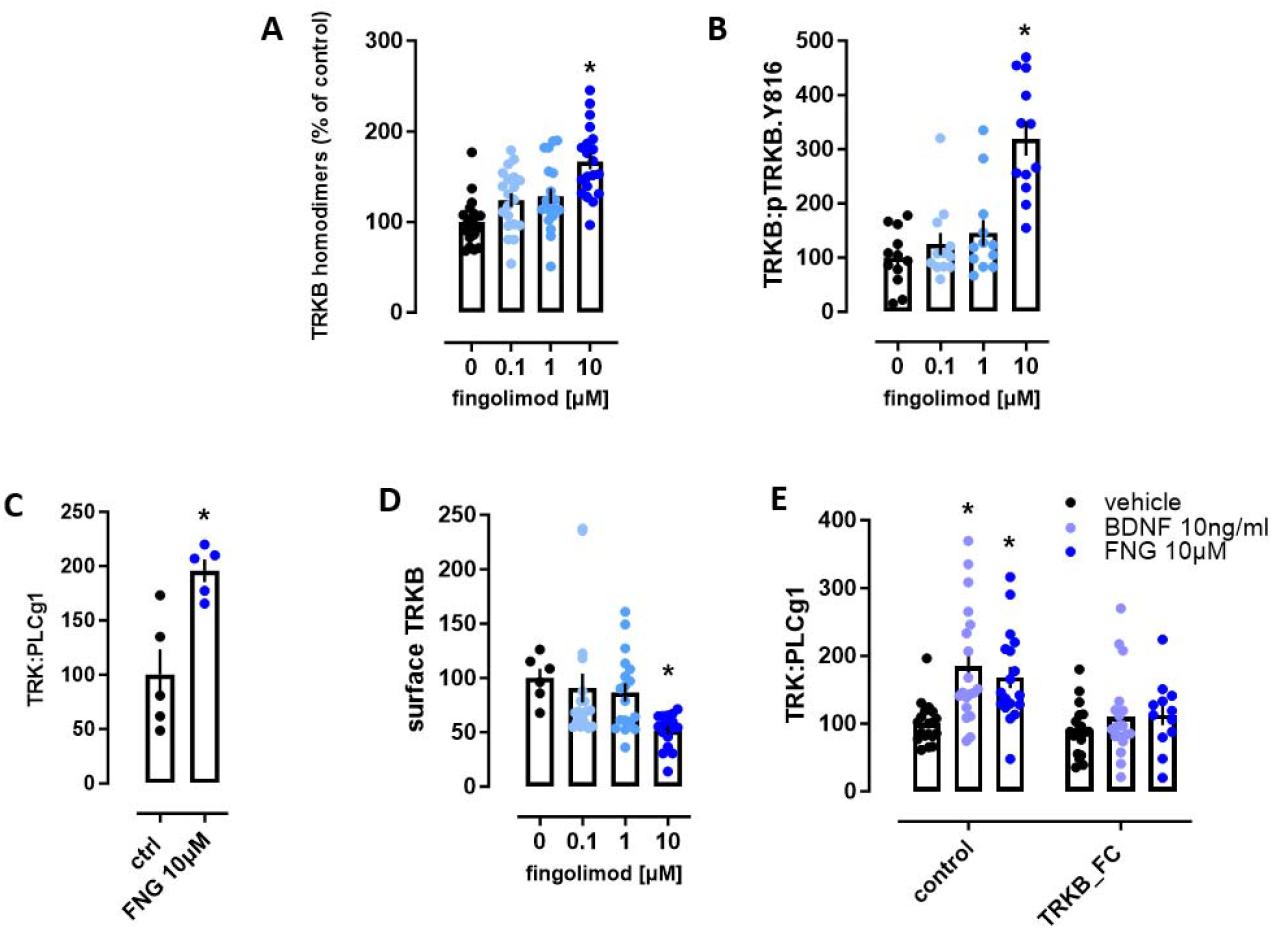
FNG induces TRKB signaling *in vitro*. (A) Acute administration of FNG (0.1, 1, 10 μM, 30 min) on N2A cells induces TRKB homodimerization (n=8/group) in PCA. Statistical analysis performed by One-way ANOVA and Tukey’s post hoc test, F (3, 76) = 13,05 p(0.1 μM)=0.0013, p(1 μM)=0.007, p(10 μM)=0.0011. (B) Acute administration of FNG and the effect on TRKB phosphorylation at tyrosine 816 measured by ELISA in primary cultured cortical cells from rat embryos (n=12/group). One-way ANOVA and Šidák’s post hoc test, F (3, 44) = 18,11, p<0.0001. (C) Acute administration of 10 μM FNG on TRKB:PLCg1 interaction measured by ELISA (n=5/group). Unpaired t-test, t=3,734, df=8, p=0.0053. (D) Detection of plasma membrane localized TRKB following acute FNG treatment (n=18/group). Kruskal-Wallis and Dunn’s post hoc test, KW= 17,81, p=0.0011. (E) Effect of the BDNF-blocking peptide TRKB_FC on the effects of acute FNG treatment in TRKB-PLCg1 interaction measured by ELISA (n=18/group). Two-way ANOVA and Tukey’s post hoc test, F (1, 100) = 15,93 p=0.0001 for the treatment effect. Data is expressed as mean+SEM.

Next, we assessed whether FNG treatment affects TRKB localization on the plasma membrane on primary cortical cells. While the lowest concentration of FNG did not alter surface TRKB levels, 10 μM FNG significantly reduced the amount of TRKB receptor at the plasma membrane at 30 min after FNG stimulation (Fig. 1D). This is consistent with the behavior of the receptor which, upon activation, is internalized and sequestered from the cell surface (Harrington and Ginty 2013; Sommerfeld et al. 2000), further indicating that FNG promotes TRKB activity and signaling. We then investigated the role of BDNF on FNG-induced modulation of TRKB activity. We found that the interaction between TRKB and its signaling partner PLCg1 induced by FNG after 30 min treatment, was blocked when primary cortical neurons were pretreated with TRKB_FC, a TRKB-like peptide that sequesters extracellular BDNF impeding its binding to TRKB receptors (Fig. 1E), indicating that BDNF is required for the effects of FNG on TRKB signaling.

Because TRKB can be transactivated by other factors in absence of BDNF, in particular by Src family kinases (Y. Z. Huang and McNamara 2010; Rajagopal and Chao 2006), we assessed their involvement in the activation of TRKB following FNG treatment. Indeed, phosphorylation at Y416, an indication of Src kinases activation, was significantly increased after 30 min treatment with FNG (Fig. 2A). However, when we treated cells with the kinase inhibitor PP1 to inhibit Src family kinases (Hanke et al. 1996), we found that FNG was still able to promote TRKB dimerization, indicating that TRKB dimerization lays upstream of other cellular players that could contribute to TRKB activation and rather initiates a positive feedback loop of co-transactivation that is likely stimulated by BDNF binding to TRKB (Zhang et al. 2013) (Fig. 2B). Then, we performed a binding assay to investigate whether FNG treatment might promote the physiological interaction between BDNF and TRKB in cortical neuronal lysates. While we could detect an expected increase in TRKB binding with increasing concentration of BDNF, FNG treatment did not modify this binding at any concentration (Fig. 2C). This suggests that while BDNF is required for TRKB activation in response to FNG, the increased TRKB activity observed following FNG treatment does not occur by direct modulation of the binding between the receptor and its ligand.

**Figure 2.**
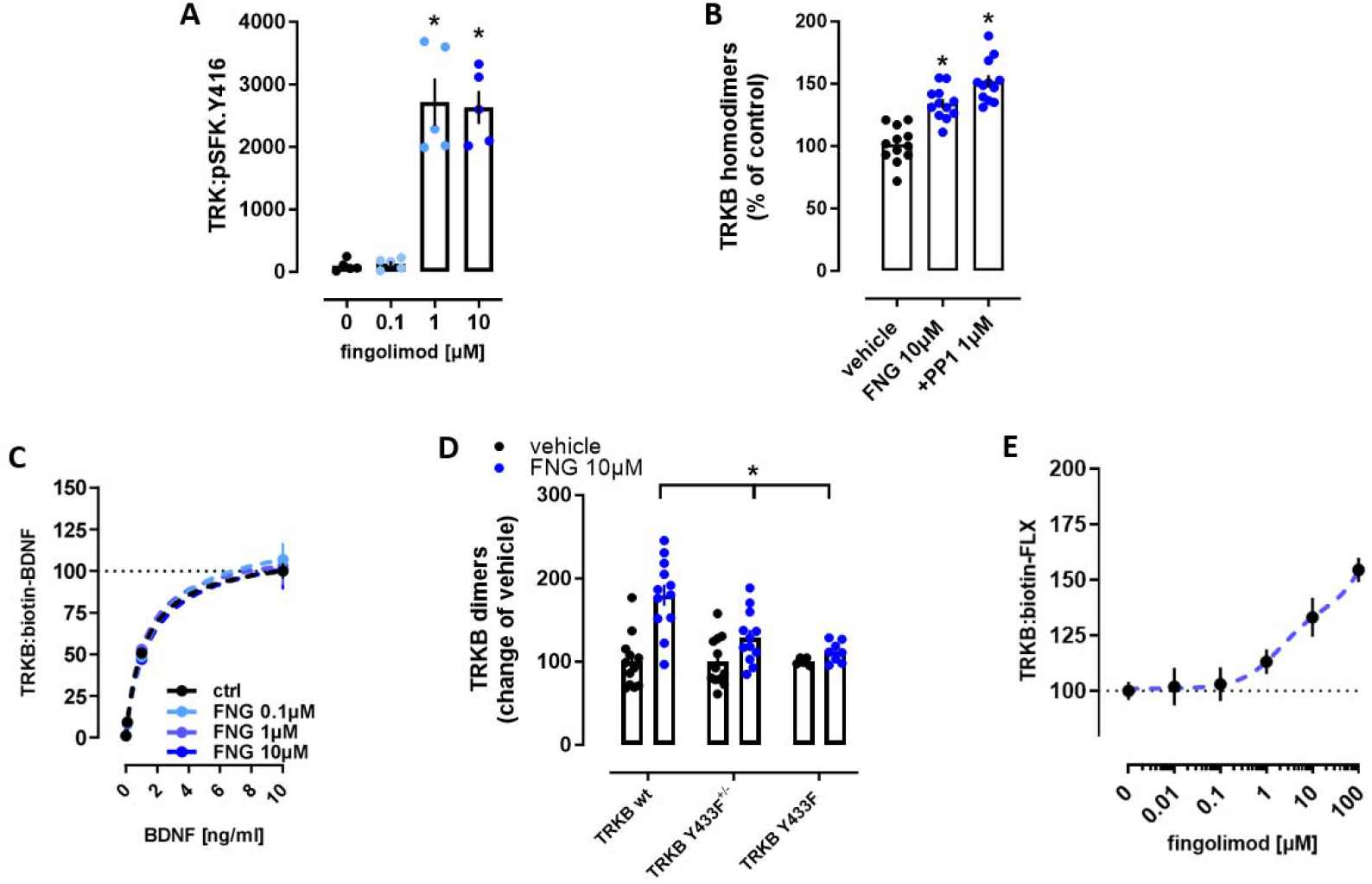
TRKB activation by FNG is an early event in FNG action and requires BDNF. (A) Acute administration of FNG and the activation of Src family kinase measured by ELISA on primary rat cortical neurons (n=5/group). One way ANOVA and Fisher’s LSD, F (3, 16) = 40,13, p(1 μM, 10 μM)<0.0001. (B) Effect of acute FNG treatments on TKRB dimerization in N2A cells in presence of kinase inhibitors PP1 and k252a (n=12/group). One way ANOVA and Tuckey’s post hoc test, F (5, 66) = 19,72, p<0.0001. (C) Ligand binding assay for TRKB and biotinylated BDNF in presence of increasing dose of FNG (n= 6/group). (D) Acute administration of FNG does not induce dimerization of the TRKB mutant Y433F (n=12/group). Two-way ANOVA and Šidák’s post hoc test, F (2, 55) = 6,216, p(Y433F^+/-^)=0.0032, p(Y433F)=0.0002. Data is expressed as mean+SEM. (E) Effect of increasing concentration of FNG on the binding of biotinylated fluoxetine on TRKB (n=6/group).

TRKB contains in its transmembrane region an inverted cholesterol-recognition domain (CARC) that mediates its interaction with membrane cholesterol (Cannarozzo et al. 2021). We have previously observed that different classes of antidepressants, including psychedelics, can directly bind to the CARC domain of TRKB and promote its activity through BDNF (Casarotto et al. 2021; Cannarozzo et al. 2021; Moliner et al. 2023). We tested the dimerization induced by FNG on wild-type TRKB, and on a mutant TRKB (Y433F) that has been shown to be insensible to cholesterol binding (Casarotto et al. 2021) (Fig. 2D). Indeed, FNG induced dimerization of wild type TRKB after 30 min of exposure, but this dimerization was reduced both for wild-type/mutant couples and between two mutant molecules, indicating that the CARC domain and the transmembrane domain of TRKB are important for FNG-induced activity. We have previously shown that cholesterol promotes the binding of the antidepressant fluoxetine to TRKB, facilitating BDNF signaling, whereas antidepressants, which interact with TRKB dimers through the same domain as fluoxetine, compete with and thereby inhibit fluoxetine binding to TRKB (Casarotto et al. 2021). We therefore tested if FNG, due to its structural properties, might modulate TRKB activity by interacting with the CARC domain. Indeed, FNG increased fluoxetine binding to TRKB in a dose-dependent way (Fig 2E), mimicking the effects of cholesterol but not that of antidepressant drugs (Casarotto et al. 2021).

### FNG compensates for the reduction of BDNF in mice exposed to stress in contextual fear conditioning

We next investigated the role of FNG in facilitating TRKB activity and whether FNG can mimic the effects of cholesterol *in vivo*. We tested whether FNG injections in wild type mice and mice lacking one allele of the Bdnf gene (BDNF.het) affected behavior in a contextual fear conditioning paradigm, with and without pravastatin pretreatment to deplete endogenous cholesterol. Cholesterol depletion by pravastatin (10mg/kg, 14 days in drinking water) has been shown to affect TRKB signaling and downstream neuroplasticity *in vitro* (Suzuki et al, 2004) and *in vivo* (Casarotto et al, 2021). In line with previous evidence, in our experimental conditions, pravastatin worsen the ability of the mice to discriminate between two separate contextual environments (Fig. 3A). Both genotypes acquired conditioning, although BDNF.het mice showed elevated freezing. However, acute injections of FNG (1 mg/kg) two hours before being exposed to foot-shocks did not influence fear conditioning (context A) in either genotype (Fig. 3B). When exposed to a novel environment (context B) 24h later, BDNF.het mice showed an elevated level of freezing, suggesting generalization of the fear response (Fig. 3C). FNG given 2h before testing did not have any effects on wild type mice while it reduced the elevated freezing response in BDNF.het animals to a level seen in the wild-type mice. When the mice were returned to context A without being exposed to foot-shocks (extinction retrieval), BDNF.het mice again showed increased freezing when compared to wild type mice, but FNG treatment did not influence freezing in either genotype (Fig. 3D). These data suggest that while FNG has no effects in a physiological context after acute injection, it sensitizes TRKB signaling to BDNF in presence of reduced levels of BDNF. In pravastatin-treated animals, FNG treatment, although not statistically significantly, has a tendency to reduce freezing in the wild-type group while it did not rescue the freezing in BDNF.het animals (Fig. 3F). This suggests that FNG effects of BDNF sensitization on TRKB may partially mimic the effects of cholesterol, explaining why wild type animals with normal amount of brain cholesterol do not respond to FNG treatment. The fact that FNG could not completely rescue the reduction of cholesterol levels indicates that the effect of FNG and cholesterol on synaptic plasticity are not fully interchangeable, and cholesterol is still required for FNG action.

**Figure 3.**
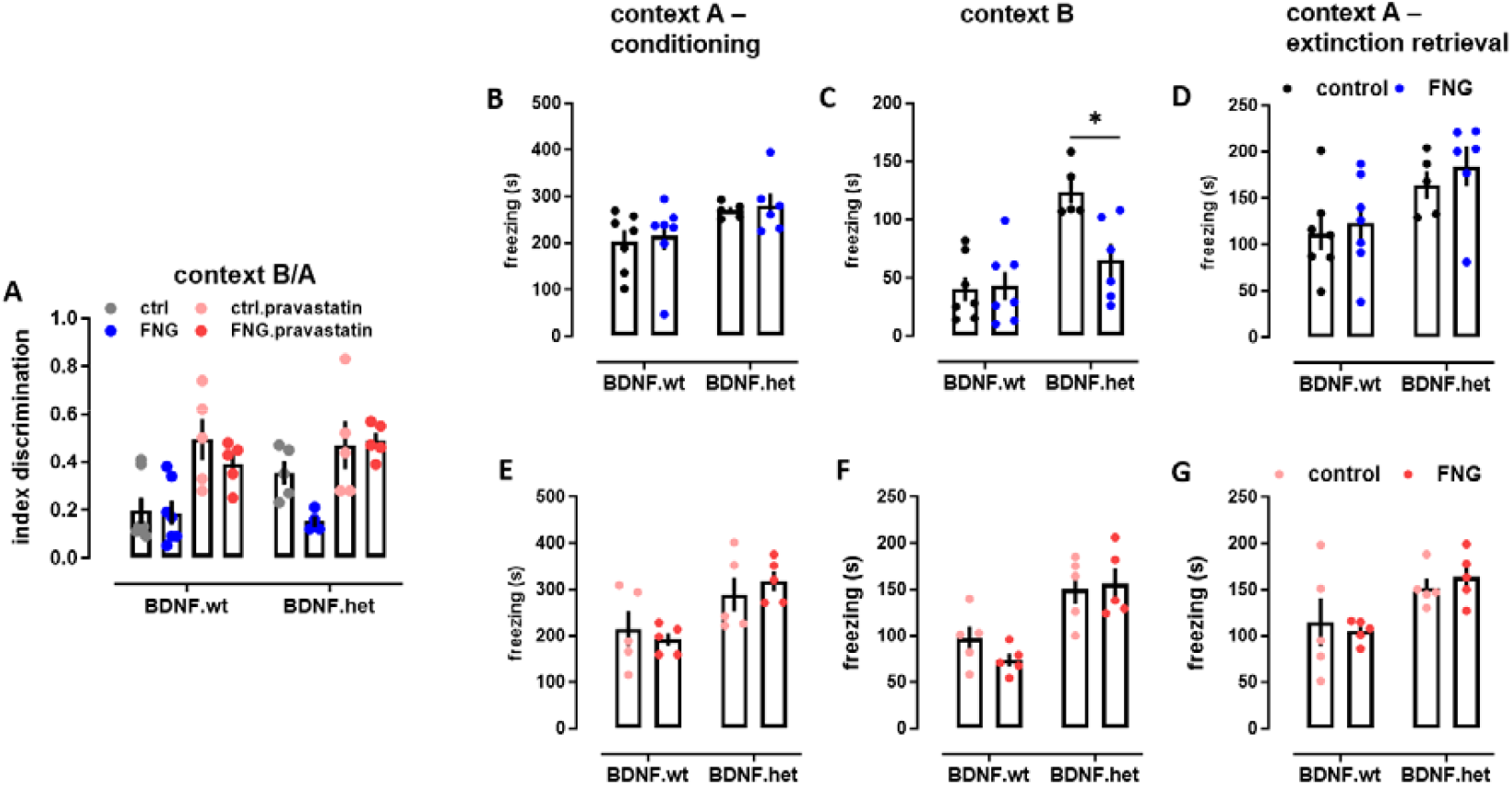
The effect of FNG administration on contextual fear conditioning. (A) Combined effects of pravastatin (10mg/kg for 14 days in drinking water) and fingolimod (1mg/kg, 2h prior) on the ability of female mice to discriminate between context A and context B, expressed as discrimination index. Pravastatin treatment impairs the discrimination ability of the two contexts, Three-way ANOVA F(1, 35) = 30,66, p<0.0001. (B) 10 min foot shock conditioning session for BDNF.wt and BDNF.het mice. FNG treatment (1 mg/kg, 2h prior) did not affect the freezing behaviour. Two way ANOVA, F(1, 21) = 9,068, p=0,9925, while BDNF.hets generally freeze more compared to control, Two way ANOVA F(1, 21)=12.78, p=0.0018. (C) FNG treatment (1 mg/kg, 2h prior, blue bars) had no effect on BDNF.wt animals exposed for 5 min to a novel environment but could rescue the generalized freezing of the BDNF.het animals. F(1, 21) = 6,279, p(interaction)=0,0205, p(controls)=0.0001, p(BDNF.het)=0.004283 by two-way ANOVA and uncorrected Fisher’s LSD multiple comparisons (D) FNG treatment did not have any effect on the freezing behavior of BDNF.wt and BDNF.het mice placed back in the conditioning environment without foot shocks. Two way ANOVA, F (1, 21) = 0,05193, p=0,8219. Similarly as the conditioning, BDNF.het freeze more compared to wild-type littermates F(1, 21)=8.636, p= p=0.0078. In a cohort of animals pre-treated with pravastatin (10mg/kg for 14 days in drinking water) (Panels E,F,G) FNG (1mg/kg, 2h prior) did not affect the conditioning. BDNF.het mice freeze significantly more compared to control animals (E), Two way ANOVA F(1,16)=11.8, p=0.0034. (F) Similarly as for the condition, BDNF.het mice freeze more than the wt littermates, Two way ANOVA, F(1,16)=24.49; p=0.0001. FNG treatment (1mg/kg, 2h prior) could not rescue the freezing behaviour in BDNF.het mice and had no significant effects on BDNF.wt animals. (G) Animals placed back in the conditioning environment without foot shock did not show any behavioral effect after FNG treatment (1mg/kg, 2h prior). As in the previous conditions, BDNF.het mice generally freeze more than the wt, Two way ANOVA, F(1,16)=9.374, p=0.0075. Data expressed as mean+SEM, n=5/7 per group.

## DISCUSSION

In this study, we identified an early mechanism for FNG, a sphingosine analog already approved for the treatment of MS, in modulating TRKB activity both *in vitro* and *in vivo*. Increased BDNF expression has been implicated in the plasticity-promoting effects of FNG (Deogracias et al. 2012; Doi et al. 2013; Golan et al. 2019; Vidal-Martinez et al. 2019; Yu et al. 2023). However, it has been unclear how this effect is mediated. BDNF mRNA is regulated by neuronal activity and calcium entry is a central stimulus for the increased BDNF expression (Zafra et al. 1990). However, activation of TRKB receptors increases BDNF mRNA production in a positive feedback loop manner (Tuvikene et al. 2016; Yasuda et al. 2007). Previous studies have shown that ERK phosphorylation is increased in response to FNG stimulation within 30 min, whereas BDNF mRNA is increased after 60 minutes (Deogracias et al. 2012), indicating that TRKB activation may contribute to the increase in BDNF mRNA levels after FNG treatment.

Here we show that acute FNG treatment of neuronal cell lines and primary cortical neurons promotes TRKB activity *in vitro* within 30 minutes. Because there is no known mechanism for FNG to directly activate TRKB, and because the short treatment time may not allow for an increase in BDNF production, we hypothesized that FNG may facilitate BDNF binding to TRKB, acting as a positive allosteric modulator, and therefore promoting further activation, similarly to what was shown previously for other plasticity-inducing drugs (Casarotto et al. 2021; Moliner et al. 2023). Indeed, FNG-induced TRKB activation was BDNF dependent, indicating that FNG acts as an allosteric potentiator of TRKB, sensitizing it to the effects of BDNF.

Sphingosine (the physiological FNG analog) is constitutively present in the plasma membrane as a metabolite of sphingomyelin and ceramide and is known to induce membrane rigidity and order, similar to cholesterol (Alanko et al. 2005; Contreras et al. 2006). Together with other sphingolipids, it associates with cholesterol and is enriched in lipid rafts and synaptic membranes (Ikonen 2008). Moreover, chronic FNG treatment alters the lipid composition of the plasma membrane in the hippocampus of healthy young mice and accumulates in lipid-rich areas of the brain (Magalhães et al. 2024). This implies that FNG can insert into the lipid bilayer due to its apolar properties and induce the formation of ordered membrane microdomains, which are also known to be signaling hubs for TRKB (Pereira and Chao 2007; Suzuki et al. 2004). Investigating whether FNG modulates TRKB by altering membrane dynamics, we found that increased dimerization is prevented in TRKB (Y433F) mutants, suggesting that the effects of FNG are targeted to the transmembrane region of TRKB (Casarotto et al. 2021). FNG did not compete for TRKB interaction with the antidepressant fluoxetine, indicating that it does not bind to the antidepressant binding site within the TRKB dimer. In contrast, FNG promoted the binding of fluoxetine to TRKB in a manner that resembles the effects of cholesterol (Casarotto et al. 2021).

Our data suggest that FNG associates with the transmembrane domain of TRKB, thereby sensitizing TRKB to BDNF, which could then lead to increased production of BDNF mRNA and protein in a positive feedback fashion, as it was previously shown (Deogracias et al, 2012). *In vivo*, FNG could rescue the behavioral effects of decreased BDNF levels in contextual fear conditioning, a behavioral paradigm to assess hippocampal synaptic plasticity and memory (Phillips and LeDoux 1992). However, in mice with reduced overall cholesterol, which are already impaired in the tasks compared to controls, FNG is not able to rescue the behavioral effects anymore, indicating that FNG and cholesterol are not mutually interchangeable and physiological levels of cholesterol are still required for an effective FNG response. Observations by other groups of FNG effects on wild-type animals in similar behavioral paradigms could be explained by the long term administration of the drug (Efstathopoulos et al. 2015; Sun et al. 2016), remaining in line with the core findings of our study. Our data indicate that FNG administration sensitizes TRKB to the effect of BDNF early on both *in vitro* and *in vivo*, which may at least in part promote the increase in BDNF production observed the day after FNG treatment.

Thus, we have identified an early action of FNG on TRKB that at least partially leads to the increased BDNF-mediated neuronal plasticity that is thought to underline many of the effects of FNG on neurological disorders.

## ACKNOWLEDGEMENTS

The authors thank Sulo Kolehmainen, Seija Lågas and the mouse behavioral phenotyping facility (MBPF, University of Helsinki) for technical assistance and Prof. Michnick and Dr. Huttunen for the original GLuc plasmids. The whole Castrén lab is thanked for insightful discussions. The work was supported by the Research Council of Finland (projects 294710, 307416, 1333589).

## CONFLICT OF INTEREST STATEMENT

EC is a co-founder and board member of Kasvu Therapeutics. PC is employed by Cell Press. The other authors claim no conflict of interest.

## AUTHOR CONTRIBUTION

CAB, PC and EC designed the experiments. CAB, JA, NS, KK, CB and PC performed the experiments. CAB and EC wrote the manuscript.

---

**Supplementary figure 1.**
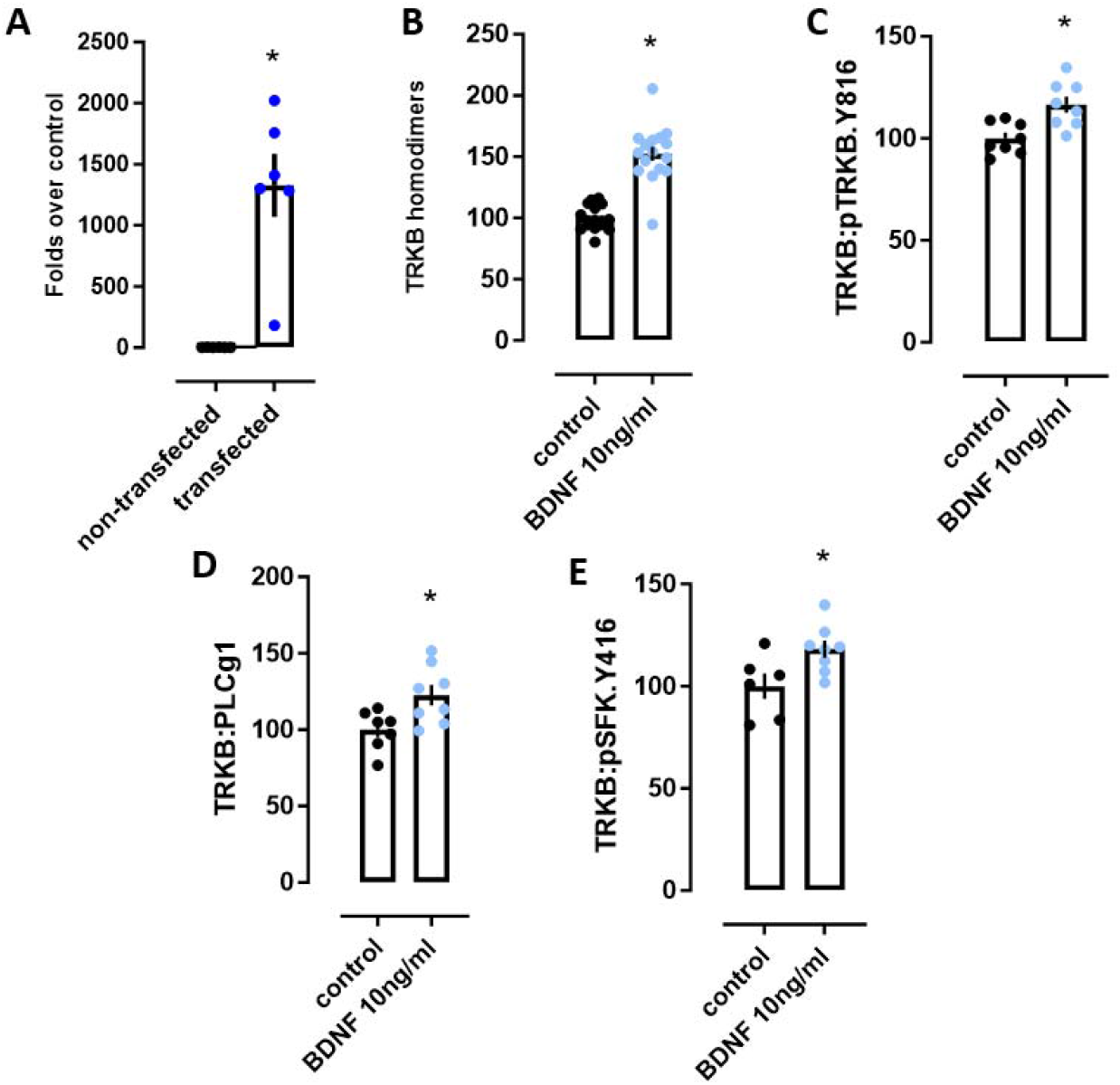
TRKB signaling *in vitro*, related to Figure 1. (A) Quantification of TRKB mRNA levels by qPCR in N2A cells non-transfected and transfected with TRKB.wt plasmid. (B) 30min BDNF treatment (10ng/ml) increase TRKB homodimerization by PCA in N2A cells. Unpaired t-test, t=8,351, df=30, p<0.0001, n=16. (C) BDNF (10ng/ml) increased TRKB phosphorylation at tyrosine 816 in cortical neurons in ELISA. Unpaired t-test, t=3,433, df=14, p=0.004, n=8. (D) BDNF (10ng/ml) increased interaction in cortical neurons between TRKB and its signaling partner PLCgamma1 in ELISA. Unpaired t-test, t=2,646, df=13, p=0.02, n=7/8. (E) BDNF (10ng/ml) promotes interaction between TRKB and activated Src family kinase in cortical neurons measured by ELISA. Unpaired t-test, t=2,512, df=12, p=0.027, n=6/8. Data is expressed as mean+SEM.

## REFERENCES

Alanko, S. M., K. K. Halling, S. Maunula, J. P. Slotte, and B. Ramstedt. 2005. “Displacement of Sterols from Sterol/Sphingomyelin Domains in Fluid Bilayer Membranes by Competing Molecules.” Biochim Biophys Acta 1715(2): 111–21. doi:10.1016/j.bbamem.2005.08.002.

Brunello, C. A., X. Yan, and H. J. Huttunen. 2016. “Internalized Tau Sensitizes Cells to Stress by Promoting Formation and Stability of Stress Granules.” Sci Rep 6: 30498. doi:10.1038/srep30498.

Buchman, A. S., L. Yu, P. A. Boyle, J. A. Schneider, P. L. De Jager, and D. A. Bennett. 2016. “Higher Brain BDNF Gene Expression Is Associated with Slower Cognitive Decline in Older Adults.” Neurology 86(8): 735–41. doi:10.1212/WNL.0000000000002387.

Cannarozzo, C., S. M. Fred, M. Girych, C. Biojone, G. Enkavi, T. Róg, I. Vattulainen, P. C. Casarotto, and E. Castrén. 2021. “Cholesterol-Recognition Motifs in the Transmembrane Domain of the Tyrosine Kinase Receptor Family: The Case of TRKB.” Eur J Neurosci 53(10): 3311–22. doi:10.1111/ejn.15218.

Casarotto, P. C., M. Girych, S. M. Fred, V. Kovaleva, R. Moliner, G. Enkavi, C. Biojone, et al. 2021. “Antidepressant Drugs Act by Directly Binding to TRKB Neurotrophin Receptors.” Cell 184(5): 1299-1313.e19. doi:10.1016/j.cell.2021.01.034.

Castrén, Eero, and Lisa M. Monteggia. 2021. “Brain-Derived Neurotrophic Factor Signaling in Depression and Antidepressant Action.” Biological Psychiatry 90(2): 128–36. doi:10.1016/j.biopsych.2021.05.008.

Chen, B., D. Dowlatshahi, G. M. MacQueen, J. F. Wang, and L. T. Young. 2001. “Increased Hippocampal BDNF Immunoreactivity in Subjects Treated with Antidepressant Medication.” Biol Psychiatry 50(4): 260–65. doi:10.1016/s0006-3223(01)01083-6.

Chen, Hui, Marc Lombès, and Damien Le Menuet. 2017. “Glucocorticoid Receptor Represses Brain-Derived Neurotrophic Factor Expression in Neuron-like Cells.” Molecular Brain 10: 12. doi:10.1186/s13041-017-0295-x.

Contreras, F. X., J. Sot, A. Alonso, and F. M. Goñi. 2006. “Sphingosine Increases the Permeability of Model and Cell Membranes.” Biophys J 90(11): 4085–92. doi:10.1529/biophysj.105.076471.

Deogracias, Rubén, Morteza Yazdani, Martijn P. J. Dekkers, Jacky Guy, Mihai Constantin S. Ionescu, Kaspar E. Vogt, and Yves-Alain Barde. 2012. “Fingolimod, a Sphingosine-1 Phosphate Receptor Modulator, Increases BDNF Levels and Improves Symptoms of a Mouse Model of Rett Syndrome.” Proceedings of the National Academy of Sciences 109(35): 14230–35. doi:10.1073/pnas.1206093109.

Deyama, S., and R. S. Duman. 2020. “Neurotrophic Mechanisms Underlying the Rapid and Sustained Antidepressant Actions of Ketamine.” Pharmacol Biochem Behav 188: 172837. doi:10.1016/j.pbb.2019.172837.

Di Pardo, A., E. Amico, M. Favellato, R. Castrataro, S. Fucile, F. Squitieri, and V. Maglione. 2014. “FTY720 (Fingolimod) Is a Neuroprotective and Disease-Modifying Agent in Cellular and Mouse Models of Huntington Disease.” Hum Mol Genet 23(9): 2251–65. doi:10.1093/hmg/ddt615.

Doi, Y., H. Takeuchi, H. Horiuchi, T. Hanyu, J. Kawanokuchi, S. Jin, B. Parajuli, et al. 2013. “Fingolimod Phosphate Attenuates Oligomeric Amyloid β-Induced Neurotoxicity via Increased Brain-Derived Neurotrophic Factor Expression in Neurons.” PLoS One 8(4): e61988. doi:10.1371/journal.pone.0061988.

Dou, S. H., Y. Cui, S. M. Huang, and B. Zhang. 2022. “The Role of Brain-Derived Neurotrophic Factor Signaling in Central Nervous System Disease Pathogenesis.” Front Hum Neurosci 16: 924155. doi:10.3389/fnhum.2022.924155.

Duman, Ronald S., and Lisa M. Monteggia. 2006. “A Neurotrophic Model for Stress-Related Mood Disorders.” Biological Psychiatry 59(12): 1116–27. doi:10.1016/j.biopsych.2006.02.013.

Dwivedi, Y., H. S. Rizavi, R. R. Conley, R. C. Roberts, C. A. Tamminga, and G. N. Pandey. 2003. “Altered Gene Expression of Brain-Derived Neurotrophic Factor and Receptor Tyrosine Kinase B in Postmortem Brain of Suicide Subjects.” Arch Gen Psychiatry 60(8): 804–15. doi:10.1001/archpsyc.60.8.804.

Efstathopoulos, P., A. Kourgiantaki, K. Karali, K. Sidiropoulou, A. N. Margioris, A. Gravanis, and I. Charalampopoulos. 2015. “Fingolimod Induces Neurogenesis in Adult Mouse Hippocampus and Improves Contextual Fear Memory.” Transl Psychiatry 5: e685. doi:10.1038/tp.2015.179.

Fred, Senem Merve, Liina Laukkanen, Cecilia A. Brunello, Liisa Vesa, Helka Göös, Iseline Cardon, Rafael Moliner, et al. 2019. “Pharmacologically Diverse Antidepressants Facilitate TRKB Receptor Activation by Disrupting Its Interaction with the Endocytic Adaptor Complex AP-2.” The Journal of Biological Chemistry 294(48): 18150–61. doi:10.1074/jbc.RA119.008837.

Golan, M., K. Mausner-Fainberg, B. Ibrahim, M. Benhamou, A. Wilf-Yarkoni, H. Kolb, K. Regev, and A. Karni. 2019. “Fingolimod Increases Brain-Derived Neurotrophic Factor Level Secretion from Circulating T Cells of Patients with Multiple Sclerosis.” CNS Drugs 33(12): 1229–37. doi:10.1007/s40263-019-00675-7.

Haapasalo, A., T. Saarelainen, M. Moshnyakov, U. Arumäe, T. R. Kiema, M. Saarma, G. Wong, and E. Castrén. 1999. “Expression of the Naturally Occurring Truncated trkB Neurotrophin Receptor Induces Outgrowth of Filopodia and Processes in Neuroblastoma Cells.” Oncogene 18(6): 1285–96. doi:10.1038/sj.onc.1202401.

Hanke, Jeffrey H., Joseph P. Gardner, Robert L. Dow, Paul S. Changelian, William H. Brissette, Elora J. Weringer, Brian A. Pollok, and Patricia A. Connelly. 1996. “Discovery of a Novel, Potent, and Src Family-Selective Tyrosine Kinase Inhibitor: STUDY OF Lck-AND FynT-DEPENDENT T CELL ACTIVATION (*).” Journal of Biological Chemistry 271(2): 695–701. doi:10.1074/jbc.271.2.695.

Harrington, A. W., and D. D. Ginty. 2013. “Long-Distance Retrograde Neurotrophic Factor Signalling in Neurons.” Nat Rev Neurosci 14(3): 177–87. doi:10.1038/nrn3253.

Huang, Y., C. Huang, and W. Yun. 2019. “Peripheral BDNF/TrkB Protein Expression Is Decreased in Parkinson’s Disease but Not in Essential Tremor.” J Clin Neurosci 63: 176–81. doi:10.1016/j.jocn.2019.01.017.

Huang, Yang Z., and James O. McNamara. 2010. “Mutual Regulation of Src Family Kinases and the Neurotrophin Receptor TrkB*.” Journal of Biological Chemistry 285(11): 8207–17. doi:10.1074/jbc.M109.091041.

Ikonen, Elina. 2008. “Cellular Cholesterol Trafficking and Compartmentalization.” Nature Reviews Molecular Cell Biology 9(2): 125–38. doi:10.1038/nrm2336.

Karege, F., G. Bondolfi, N. Gervasoni, M. Schwald, J. M. Aubry, and G. Bertschy. 2005. “Low Brain-Derived Neurotrophic Factor (BDNF) Levels in Serum of Depressed Patients Probably Results from Lowered Platelet BDNF Release Unrelated to Platelet Reactivity.” Biol Psychiatry 57(9): 1068–72. doi:10.1016/j.biopsych.2005.01.008.

Karpova, N. N., A. Pickenhagen, J. Lindholm, E. Tiraboschi, N. Kulesskaya, A. Agústsdóttir, H. Antila, et al. 2011. “Fear Erasure in Mice Requires Synergy between Antidepressant Drugs and Extinction Training.” Science 334(6063): 1731–34. doi:10.1126/science.1214592.

Kartalou, G. I., A. R. Salgueiro-Pereira, T. Endres, A. Lesnikova, P. Casarotto, P. Pousinha, K. Delanoe, et al. 2020. “Anti-Inflammatory Treatment with FTY720 Starting after Onset of Symptoms Reverses Synaptic Deficits in an AD Mouse Model.” Int J Mol Sci 21(23). doi:10.3390/ijms21238957.

Laukkanen, L., C. R. A.F Diniz, S. Foulquier, J. Prickaerts, E. Castrén, and P. C. Casarotto. 2021. “Facilitation of TRKB Activation by the Angiotensin II Receptor Type-2 (AT2R) Agonist C21.” Pharmaceuticals (Basel) 14(8). doi:10.3390/ph14080773.

Magalhães, Daniela M., Nicolas A. Stewart, Myrthe Mampay, Sara O. Rolle, Chloe M. Hall, Emad Moeendarbary, Melanie S. Flint, et al. 2024. “The Sphingosine 1-Phosphate Analogue, FTY720, Modulates the Lipidomic Signature of the Mouse Hippocampus.” Journal of Neurochemistry 168(6): 1113–42. doi:10.1111/jnc.16073.

Mandala, Suzanne, Richard Hajdu, James Bergstrom, Elizabeth Quackenbush, Jenny Xie, James Milligan, Rosemary Thornton, et al. 2002. “Alteration of Lymphocyte Trafficking by Sphingosine-1-Phosphate Receptor Agonists.” Science (New York, N.Y.) 296(5566): 346–49. doi:10.1126/science.1070238.

Merezhko, M., C. A. Brunello, X. Yan, H. Vihinen, E. Jokitalo, R. L. Uronen, and H. J. Huttunen. 2018. “Secretion of Tau via an Unconventional Non-Vesicular Mechanism.” Cell Rep 25(8): 2027-2035.e4. doi:10.1016/j.celrep.2018.10.078.

Miguez, A., G. García-Díaz Barriga, V. Brito, M. Straccia, A. Giralt, S. Ginés, J. M. Canals, and J. Alberch. 2015. “Fingolimod (FTY720) Enhances Hippocampal Synaptic Plasticity and Memory in Huntington’s Disease by Preventing p75NTR up-Regulation and Astrocyte-Mediated Inflammation.” Hum Mol Genet 24(17): 4958–70. doi:10.1093/hmg/ddv218.

Moliner, Rafael, Mykhailo Girych, Cecilia A. Brunello, Vera Kovaleva, Caroline Biojone, Giray Enkavi, Lina Antenucci, et al. 2023. “Psychedelics Promote Plasticity by Directly Binding to BDNF Receptor TrkB.” Nature Neuroscience 26(6): 1032–41. doi:10.1038/s41593-023-01316-5.

Naegelin, Yvonne, Jens Kuhle, Sabine Schädelin, Alexandre N. Datta, Stefano Magon, Michael Amann, Christian Barro, et al. 2021. “Fingolimod in Children with Rett Syndrome: The FINGORETT Study.” Orphanet Journal of Rare Diseases 16(1): 19. doi:10.1186/s13023-020-01655-7.

Patnaik, A., E. Spiombi, A. Frasca, N. Landsberger, M. Zagrebelsky, and M. Korte. 2020. “Fingolimod Modulates Dendritic Architecture in a BDNF-Dependent Manner.” Int J Mol Sci 21(9). doi:10.3390/ijms21093079.

Pereira, D. B., and M. V. Chao. 2007. “The Tyrosine Kinase Fyn Determines the Localization of TrkB Receptors in Lipid Rafts.” J Neurosci 27(18): 4859–69. doi:10.1523/JNEUROSCI.4587-06.2007.

Phillips, R. G., and J. E. LeDoux. 1992. “Differential Contribution of Amygdala and Hippocampus to Cued and Contextual Fear Conditioning.” Behav Neurosci 106(2): 274–85. doi:10.1037//0735-7044.106.2.274.

Rajagopal, Rithwick, and Moses V. Chao. 2006. “A Role for Fyn in Trk Receptor Transactivation by G-Protein-Coupled Receptor Signaling.” Molecular and Cellular Neurosciences 33(1): 36–46. doi:10.1016/j.mcn.2006.06.002.

Remy, I., and S. W. Michnick. 2006. “A Highly Sensitive Protein-Protein Interaction Assay Based on Gaussia Luciferase.” Nat Methods 3(12): 977–79. doi:10.1038/nmeth979.

Smith, M. A., S. Makino, R. Kvetnanský, and R. M. Post. 1995. “Effects of Stress on Neurotrophic Factor Expression in the Rat Brain.” Annals of the New York Academy of Sciences 771: 234–39. doi:10.1111/j.1749-6632.1995.tb44684.x.

Sommerfeld, M. T., R. Schweigreiter, Y. A. Barde, and E. Hoppe. 2000. “Down-Regulation of the Neurotrophin Receptor TrkB Following Ligand Binding. Evidence for an Involvement of the Proteasome and Differential Regulation of TrkA and TrkB.” J Biol Chem 275(12): 8982–90. doi:10.1074/jbc.275.12.8982.

Sun, Y., F. Hong, L. Zhang, and L. Feng. 2016. “The Sphingosine-1-Phosphate Analogue, FTY-720, Promotes the Proliferation of Embryonic Neural Stem Cells, Enhances Hippocampal Neurogenesis and Learning and Memory Abilities in Adult Mice.” Br J Pharmacol 173(18): 2793–2807. doi:10.1111/bph.13557.

Suzuki, S., T. Numakawa, K. Shimazu, H. Koshimizu, T. Hara, H. Hatanaka, L. Mei, B. Lu, and M. Kojima. 2004. “BDNF-Induced Recruitment of TrkB Receptor into Neuronal Lipid Rafts: Roles in Synaptic Modulation.” J Cell Biol 167(6): 1205–15. doi:10.1083/jcb.200404106.

Tuvikene, J., P. Pruunsild, E. Orav, E. E. Esvald, and T. Timmusk. 2016. “AP-1 Transcription Factors Mediate BDNF-Positive Feedback Loop in Cortical Neurons.” J Neurosci 36(4): 1290–1305. doi:10.1523/JNEUROSCI.3360-15.2016.

Vidal-Martinez, G., K. Najera, J. D. Miranda, C. Gil-Tommee, B. Yang, J. Vargas-Medrano, V. Diaz-Pacheco, and R. G. Perez. 2019. “FTY720 Improves Behavior, Increases Brain Derived Neurotrophic Factor Levels and Reduces α-Synuclein Pathology in Parkinsonian GM2+/-Mice.” Neuroscience 411: 1–10. doi:10.1016/j.neuroscience.2019.05.029.

Yasuda, Makoto, Mamoru Fukuchi, Akiko Tabuchi, Masahiro Kawahara, Hiroshi Tsuneki, Yuko Azuma, Yusuke Chiba, and Masaaki Tsuda. 2007. “Robust Stimulation of TrkB Induces Delayed Increases in BDNF and Arc mRNA Expressions in Cultured Rat Cortical Neurons via Distinct Mechanisms.” Journal of Neurochemistry 103(2): 626–36. doi:10.1111/j.1471-4159.2007.04851.x.

Yu, Xueli, Xueyu Qi, Long Wei, Liansheng Zhao, Wei Deng, Wanjun Guo, Qiang Wang, et al. 2023. “Fingolimod Ameliorates Schizophrenia-like Cognitive Impairments Induced by Phencyclidine in Male Rats.” British Journal of Pharmacology 180(2): 161–73. doi:10.1111/bph.15954.

Zafra, F., B. Hengerer, J. Leibrock, H. Thoenen, and D. Lindholm. 1990. “Activity Dependent Regulation of BDNF and NGF mRNAs in the Rat Hippocampus Is Mediated by Non-NMDA Glutamate Receptors.” The EMBO journal 9(11): 3545–50. doi:10.1002/j.1460-2075.1990.tb07564.x.

Zagrebelsky, Marta, and Martin Korte. 2014. “Form Follows Function: BDNF and Its Involvement in Sculpting the Function and Structure of Synapses.” Neuropharmacology 76 Pt C: 628–38. doi:10.1016/j.neuropharm.2013.05.029.

Zhang, Zitao, Jin Fan, Yongxin Ren, Wei Zhou, and Guoyong Yin. 2013. “The Release of Glutamate from Cortical Neurons Regulated by BDNF via the TrkB/Src/PLC-Γ1 Pathway.” Journal of Cellular Biochemistry 114(1): 144–51. doi:10.1002/jcb.24311.

Zuccato, C., M. Marullo, P. Conforti, M. E. MacDonald, M. Tartari, and E. Cattaneo. 2008. “Systematic Assessment of BDNF and Its Receptor Levels in Human Cortices Affected by Huntington’s Disease.” Brain Pathol 18(2): 225–38. doi:10.1111/j.1750-3639.2007.00111.x.

